# Dynamic evolution of MLKL in primate and poxvirus genomes indicates necroptosis is a critical, not auxiliary, countermeasure during infection

**DOI:** 10.1101/2021.02.19.432003

**Authors:** Suzette Palmer, Sruthi Chappidi, Chelsea Pinkham, Dustin C. Hancks

**Author notes:** Corresponding author: (DH).

## Abstract

Pathogen infection triggers host innate defenses which can lead to the activation of regulated cell death (RCD) pathways such as apoptosis. Given a key role in immunity, apoptotic effectors are often counteracted by pathogen-encoded antagonists. Mounting evidence indicates that programmed necrosis, which is mediated by the RIPK3/MLKL axis and termed necroptosis, evolved as a countermeasure to pathogen-mediated inhibition of apoptotic signaling. However, whether this emerging inflammatory RCD pathway functions primarily as a “back-up” or fundamental response remains inconclusive. We hypothesized that if necroptosis is an instrumental defense, then its effectors should display specific signatures associated with pathogen conflict that are rare in combination: rapid evolution, viral homolog hereafter virolog, and induction by cytokines (e.g. interferons). Our rapid evolution analysis across the necroptosis pathway revealed: 1) strong signatures of positive selection for RIPK3 and MLKL in primate genomes and to a lesser extent DAI/ZBP1, 2) elevated rates of amino acid substitution on multiple surfaces including the RIPK3/MLKL binding interface and 3) evidence supporting a means of activating RIPK3 independent of homotypic RHIM domain interactions. Interestingly, a poxvirus MLKL homolog has recently been identified that acts as a RIPK3 pseudosubstrate. Our findings indicate that poxvirus MLKLs are also subject to similar but distinct volatile patterns of evolution comparable to host necroptotic factors. Specifically, viral MLKLs have undergone numerous gains and losses in poxvirus evolution with some species harboring three distinct copies. Furthermore, we confirm that MLKL can be induced by cytokines like interferon gamma. In summary, MLKL displays all three hallmarks of pivotal immune factors of which only OAS1, but not other factors like cGAS, APOBEC3G, or PKR, exhibits. These data support the hypothesis that over evolutionary time, necroptosis has served as a key battleground during infection and is therefore, not an auxiliary response.

**Summary:** Regulated cell death (RCD), such as apoptosis, is a common host defense against invading pathogens. Necroptosis, an inflammatory RCD pathway, is thought to have emerged as an auxiliary response when other cell death pathways are suppressed by pathogens during infection. In our analyses, we have identified genetic changes in host and viral factors associated with necroptosis that display signatures of adaptation and may have served as evolutionary countermeasures to shape infection outcomes. Consistent with repeated targeting by pathogen-encoded inhibitors, we found robust signatures of rapid evolution for the essential catalysts of necroptosis, RIPK3 and MLKL. Notably, an evolutionary signature specific to RIPK3 for a domain shared with other necroptotic factors suggests an undefined means to trigger this host defense pathway. In contrast, poxviruses appear to circumvent this pathway by constantly altering the number and nature of factors they deploy to suppress necroptosis including a mimic of MLKL, which was stolen from infected cells. Collectively, our findings provide new insights into host and viral genetics that may influence infection outcomes and the factors shaping the ability of pathogens to infect and spread to new species. Furthermore, these data support the notion that necroptosis is a fundamental, not auxiliary, host response during infection.

## Introduction

Regulated cell death (RCD) programs are pivotal host defense responses against pathogens [1–3]. Consequently, pathogens have evolved a myriad of strategies to counteract these responses [4, 5], which includes the activation of apoptosis. Apoptosis is a well-characterized, non-inflammatory form of RCD that was first classified by its distinct morphology that includes membrane blebbing, apoptotic bodies, and nuclear and cytoplasmic condensation [6]. Both extrinsic as well as intrinsic cues can trigger a signaling cascade that involves the activation of effector caspases to execute apoptosis (Fig 1A) [7] which was characterized, in large part, using viral proteins [8].

**Fig 1.**
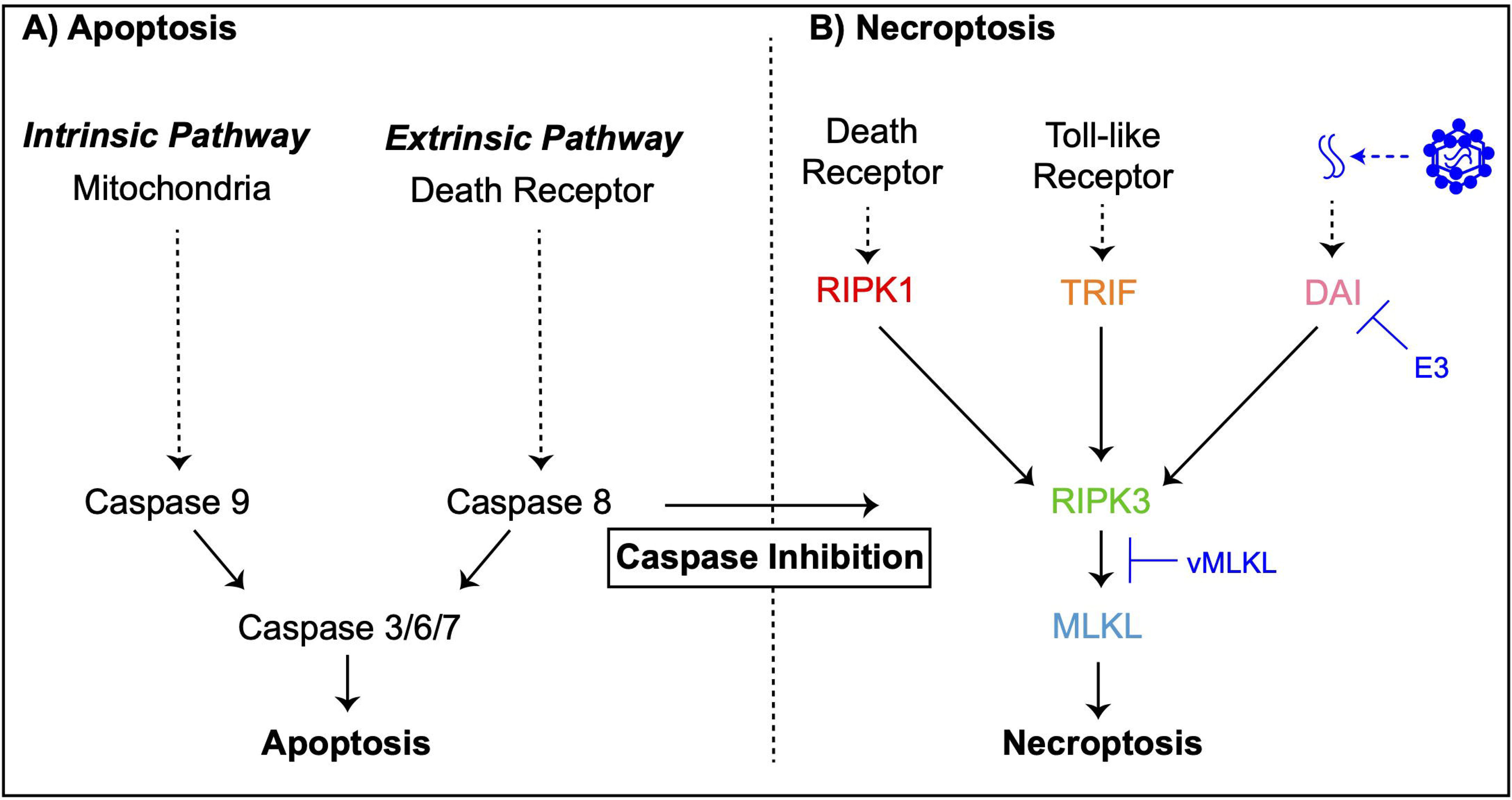
Cell death pathways, a mechanism of host defense. **(A)** Apoptosis is a regulated cell death pathway that acts as a key line of host defense against pathogens and is executed by caspases via an extrinsic or intrinsic route [7]. **(B)** Necroptosis can act as an alternative regulated cell death pathway in the event of inhibition of the apoptosis pathway by pathogens [10] such as poxviruses. E3 [32] and vMLKL [31] are recently described poxvirus factors that inhibit necroptotic signaling.

Recently, a RCD pathway defined as programmed necrosis, hereafter necroptosis, was discovered. Necroptosis is triggered when specific apoptotic signaling and effector functions are suppressed during pathogen infection. Given the conditions under which it is activated, necroptosis has often been considered a necessary “back-up” host response to pathogen-mediated inhibition of apoptosis (Fig 1B) [9, 10]. In contrast to apoptotic cell death, necroptosis is inflammatory and characterized by cell and organelle swelling followed by plasma membrane rupture [7, 11, 12]. Apoptotic and necroptotic signaling also differ in their evolutionary dynamics over large timescales. Specifically, key effectors of cellular necroptotic signaling emerged later in evolution and display a “patchy” phylogenetic distribution [13, 14] which is in marked contrast to the more ancient and highly conserved components of apoptotic signaling [15–17]. Notably, the evolutionary dynamics of necroptotic signaling over more recent timescales is unknown but may inform determinants shaping contemporary infections.

Necroptosis can be initiated by at least three presumably independent receptors: <u>d</u>eath receptor (DR) signaling, 2) the <u>p</u>athogen recognition receptor (PRR) <u>Z</u>-DNA <u>B</u>inding <u>P</u>rotein 1 (ZBP1) / <u>D</u>NA <u>A</u>ctivator of Interferon (DAI), and 3) <u>T</u>oll-like <u>R</u>eceptor 3/4 (TLR3/TLR4) [7]. Activation of DR signaling triggers <u>R</u>eceptor Interacting <u>P</u>rotein <u>K</u>inase <u>1</u> (RIPK1) to self-oligomerize, which leads to the recruitment of <u>R</u>eceptor Interacting <u>P</u>rotein <u>K</u>inase <u>3</u> (RIPK3) via physical <u>R</u>IP <u>H</u>omotypic Interaction <u>M</u>otif (RHIM) domain interactions. Activated RIPK3 subsequently binds and phosphorylates <u>M</u>ixed Lineage <u>K</u>inase-<u>L</u>ike (MLKL) [18–20]. Phosphorylation of MLKL results in a conformational change, enabling self-oligomerization of this factor to complete necroptosis through MLKL-plasma membrane destabilization [21, 22].

Importantly, DR initiation of necroptosis appears to function when caspase-8 activity is inhibited during apoptosis [23, 24]. In contrast, DAI triggers necroptosis upon binding foreign nucleic acids followed by activation of RIPK3 via RHIM-dependent interactions [25, 26]. In addition, TLR3 and TLR4 signaling can activate RIPK3 via TIR-domain-containing adaptor-inducing interferon-β (TRIF) also through physical RHIM domain interactions [27, 28]. The diverse and distinct means of activating necroptosis with convergence on the RIPK3/MLKL axis reflect the breadth of pathogens this response may protect against.

Consistently, a growing list of viral- and bacterial-encoded inhibitors that target discrete steps of necroptosis has emerged [29–36]. However, whether this pathogen-mediated antagonism has shaped cellular necroptotic factors remains unknown. Strong selective pressure may be imposed on host factors to escape direct binding by pathogen-encoded antagonists. This pressure is often visible as elevated rates of amino acid substitution, a signature of rapid evolution and a hallmark of positive selection. Signatures of rapid evolution have been observed across orthologous sequences from closely related species (e.g. primates) for several key immune factors: PKR [37], OAS1 [38, 39], MxA [40], TRIM5α [41], APOBEC3G [41], and cGAS [38, 39]. Studies of host-pathogen co-evolution may reveal novel insights into cellular responses including evidence for undefined host components, genetic determinants shaping infection outcomes, and pathogen countermeasures of these defenses. We hypothesized that analysis of signatures characteristic of host-pathogen conflict for both cellular and viral components would illuminate whether necroptosis has served as a “back-up” or a core, fundamental response over evolutionary time.

Consistent with pivotal roles in host defense across species, we report that the necroptotic axis in primates - RIPK3 and MLKL - displays widespread and recurrent signatures of rapid evolution. Interestingly, we found evidence for positive selection within the RHIM domain of RIPK3, but not at homologous sites in RHIM domains of DAI, TRIF, or RIPK1, that is proximal to a known target-site for a conserved bacterial protease. Notably, these bacterial proteases have been shown to cleave all of the RHIM domain containing necroptosis factors [30]. These data suggest an undefined means of RHIM-independent RIPK3 activation. Evolutionary genetics analysis of a recently described poxvirus MLKL homolog indicate that this antagonist has been subjected to repeated gains and losses over viral evolution with some poxvirus genomes encoding up to three mimics. In summary, our data demonstrate that MLKL belongs to a small class of key immune factors, which includes OAS1 but not PKR or cGAS, that display all three hallmarks of pivotal immune factors - rare in combination: 1) rapidly evolving, 2) viral homolog, and 3) upregulation by cytokines such as interferons. Collectively, our data are consistent with necroptosis being a fundamental host defense, not an auxiliary one, shaping infection outcomes over evolution.

## Results

### Widespread signatures of rapid evolution in primate RIPK3 and MLKL

To determine if known components of the necroptosis pathway display signatures of positive selection, we analyzed a matching set of twenty-one primate sequences for *TRIF*, *DAI*, *RIPK1*, *RIPK3* and *MLKL* using a series of evolutionary analyses. To test for recurrent positive selection, we used a combination of maximum-likelihood based algorithms, which estimate rates of non-synonymous amino acid replacements (dN) relative to synonymous (dS) amino acid substitutions. dN/dS values > 1 are considered a hallmark of positive selection. Using codon-based models implemented in <u>P</u>hylogenetic <u>A</u>nalysis by <u>M</u>aximum <u>L</u>ikelihood (PAML) [42], we detected robust gene-wide rapid evolution signatures for *MLKL* [M7 vs. M8 (F3×4) p < 5.400 x 10^- 10^], *RIPK3* [M7 vs. M8 (F3×4) p < 3.056 x 10^-9^] and to a lesser extent, *DAI* [M7 vs. M8 (F3×4) p < 5.767 x 10^-3^] but not *RIPK1* [M7 vs. M8 (F3×4) p < 0.065] or *TRIF* [M7 vs. M8 (F3×4) p < 0.066] (Fig 2A, S1 File). The signature of selection for the main effectors of the necroptosis pathway, RIPK3 and MLKL, is comparable to other key host defense factors – PKR, cGAS, OAS1 – which are also upregulated by cytokines. Indeed, MLKL is also upregulated by cytokines like interferons [43–45], which we confirm for IFNγ (Fig 2B).

**Fig 2.**
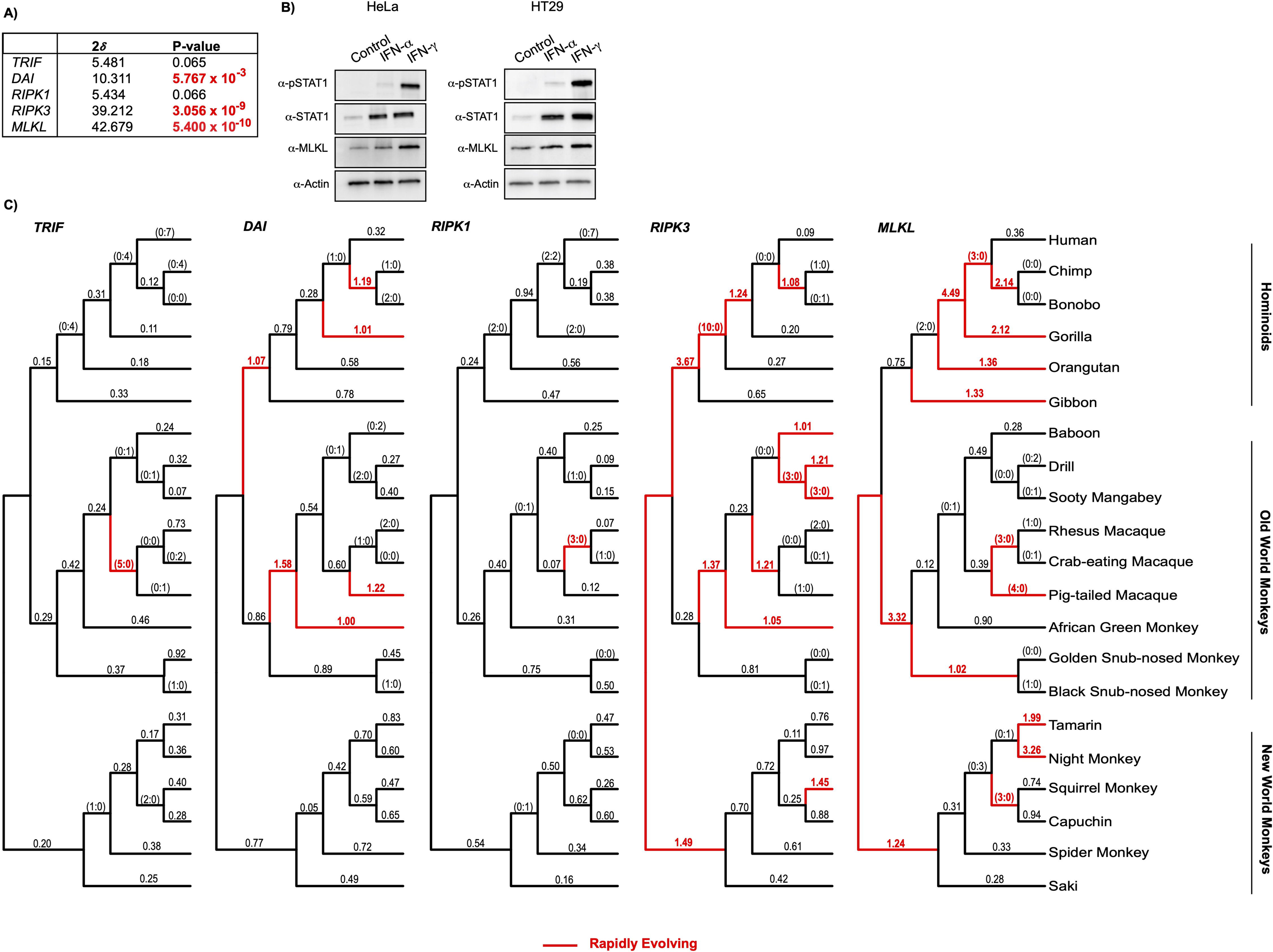
Rapid evolution of the necroptotic axis, RIPK3 and MLKL, across primate lineages. **A)** Summary of rapid evolution results for M7 x M8 (F3×4) Likelihood ratio test statistics (2*δ)* for necroptotic factors across a set of twenty-one matching primate species. **B)** MLKL is upregulated by IFNγ treatment in HeLa and HT29 cell lines. **C)** dN/dS values estimated using PAML across twenty-one primate lineages. Branches colored in red display signatures of positive selection (dN/dS > 1, or greater than or equal to 3 nonsynonymous amino acid changes relative to zero synonymous amino acid changes).

To assay when and to what extent selection pressure has shaped necroptotic signaling over primate evolution, we estimated dN/dS values across the primate phylogeny (Fig 2C) for each factor. Consistent with our gene-wide tests of evolution, we observed recurrent and widespread signatures of positive selection across primates for *RIPK3*, *MLKL* and again to a lesser extent, *DAI*. Specifically, numerous lineages across primate evolution for both *RIPK3* and *MLKL*, including branches in each major group, show substitution patterns that are characteristic of genetic conflict. Strikingly, especially strong signatures are evident for *RIPK3* in the lineage preceding the divergence of Hominidae [10 nonsynonymous changes: 0 synonymous changes] and for *MLKL* in the lineage prior to Hominidae divergence [dN/dS = 4.49] (Fig 2C). In contrast, only one lineage with a rapid evolution signature was detected for both *TRIF* and *RIPK1*. Thus, repeated innovation in the necroptosis pathway has been primarily focused on the downstream axis of *RIPK3*/*MLKL* during primate evolution.

To uncover where strong selection pressure may have been imposed within the effector proteins of necroptosis, dN/dS values were estimated for individual amino acid positions using three distinct but commonly implemented methods: PAML [42], MEME [46], and FUBAR [47] (Fig 3A, S1 File). The distribution and the number of rapidly evolving sites are thought to reflect the number of protein surfaces in genetic conflict with other factors, such as pathogen-encoded inhibitors [48]. In agreement with necroptosis being triggered by diverse pathogens, predicted rapidly evolving sites are distributed throughout the necroptosis proteins (*MLKL*, *RIPK3*, *DAI*, *RIPK1*, and *TRIF*). In support of our preceding analysis, *MLKL* (20 total sites / 11 by multiple methods) and *RIPK3* (27 total sites / 7 by multiple methods) have the most sites identified (Fig 3A). Interestingly, the majority (70%) of MLKL positively selected sites are localized to the kinase-like domain, with 12/14 sites within this domain occurring between amino acid positions 370-470 (Fig 3A). Positively selected sites mapped onto a previously published co-crystal structure of mouse RIPK3 and MLKL [49] indicate that clusters of sites represent distinct surfaces (Fig 3B). Analysis of space-filling models highlights that the majority of the rapidly evolving sites are found on the protein surface for both RIPK3 and MLKL, which suggests they represent interfaces for protein-protein interactions (S1 Fig). Notably, there is a cluster of positively selected sites for RIPK3 that surrounds MLKL and corresponds to the hydrophobic pocket where RIPK3 and MLKL interact via hydrophobic interactions (Fig 3C). This signature may indicate the presence of MLKL pseudosubstrates, as this pocket has been demonstrated previously to be necessary for RIPK3:MLKL interaction and for necroptosis to occur [31, 49].

**Fig 3.**
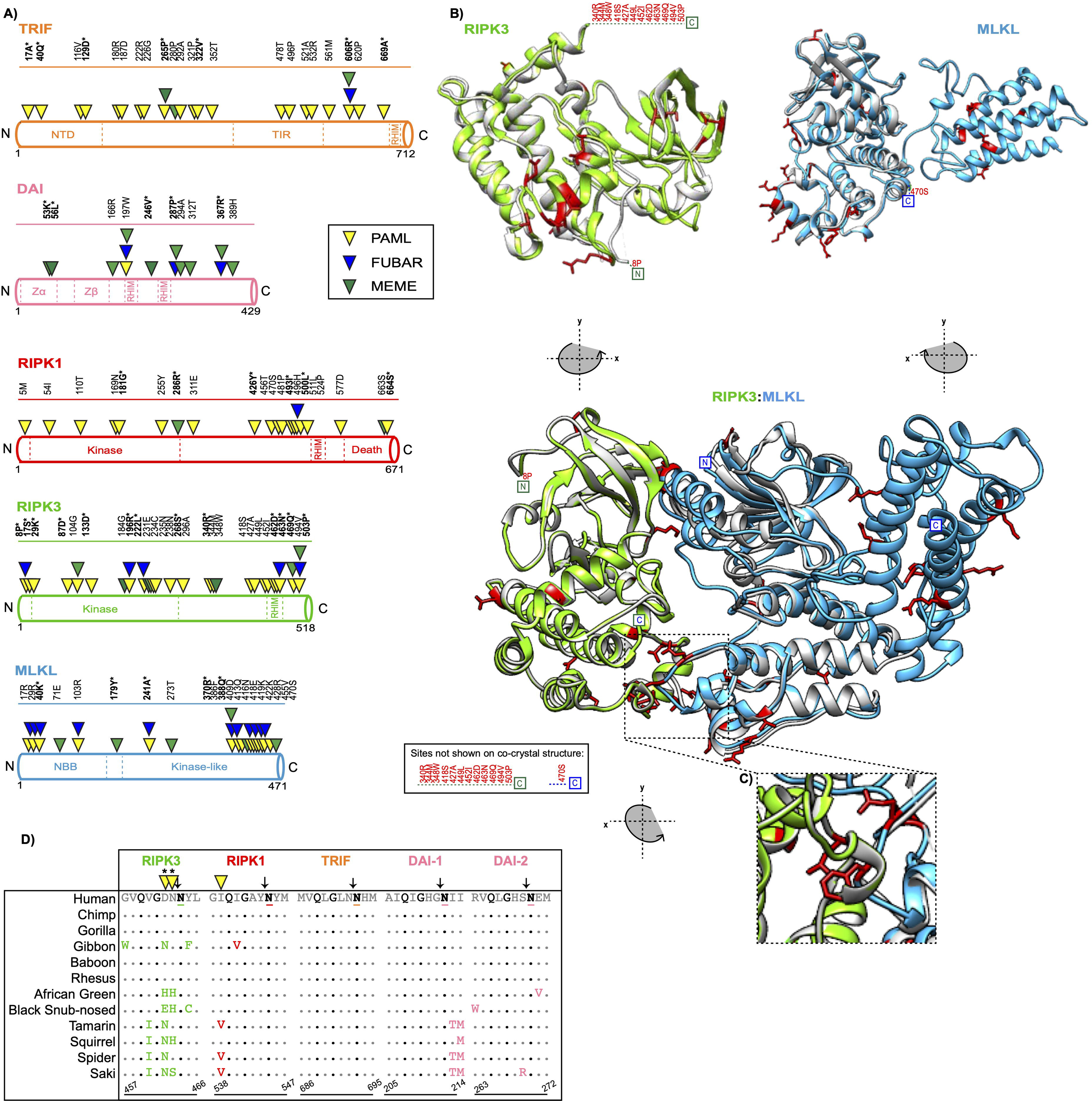
Widespread signatures of positive selection across necroptosis protein sequences. **A)** Rapidly evolving sites represented as triangles and shown for TRIF (orange), DAI (pink), RIPK1 (red), RIPK3 (green) and MLKL (blue). Rapidly evolving amino acid sites predicted using PAML (yellow triangle), FUBAR (blue triangle) and MEME (green triangle). Sites with p-values < 0.05 or posterior probabilities (P) > 0.95 are listed above the protein cartoons and represented as triangles. Sites with p-values < 0.01 or posterior probabilities (P) > 0.99 are bolded and have an asterisk. Amino acid alignment position refers to the human reference sequences. **B)** Location of rapidly evolving sites (shown in red) on the h(uman)RIPK3 (green) homology model and the hMLKL (light blue) crystal structure [20]. The homology model was predicted using Swiss-Model [64] and aligned to crystal structures m(ouse)RIPK3 (silver), mMLKL (silver), and mRIPK3:mMLKL (silver) [49]. **C)** Rapid evolution (red) at the mRIPK3 hydrophobic pocket. **D)** Rapid evolution exclusive to the RIPK3 RHIM-domain proximal to the EspL cleavage motif (Q*G**N - highlighted in black). Sequences spanning the EspL cleavage motif from 12 primates for RIPK3 (green), RIPK1 (red), TRIF (orange), and DAI (pink) RHIM domains. The EspL cleavage site is underlined and displays a black arrow above the amino acid. Sites with p-values < 0.05 or posterior probabilities (P) > 0.95 are listed above the protein cartoons and represented as triangles. Sites with p-values < 0.01 or posterior probabilities (P) > 0.99 are bolded and have an asterisk. Amino acid alignment position refers to the human reference sequences. Note: PAML, MEME, and FUBAR analysis were performed for RIPK1 without drill monkey, since this region was trimmed in previous analyses.

### Rapid evolution exclusive to the RHIM domain of RIPK3 localizes to a known bacterial protease cleavage site

RHIM-RHIM homotypic interactions between RIPK3 and established necroptosis activators RIPK1, TRIF, and DAI are required for signal transduction. The critical nature of this interaction is also supported by the identification of both viral- and bacterial- encoded inhibitors that target the RHIM domains of these proteins. These include cytomegalovirus M45 and herpes simplex virus ICP6 and ICP10, which all encode RHIM domains and target host RHIM-dependent interactions [29, 33]. Another example is EspL, a bacterial protease, which is encoded by diverse species, including enteropathogenic *Escherichia coli* (EPEC), *Salmonella enterica,* and *Yersinia pestis* [30]. Notably, EPEC EspL has been shown to cleave a specific motif (Q*G**N) in the RHIM domains of human and mouse TRIF, DAI, RIPK3, and RIPK1, which results in the suppression of necroptosis [30]. Interestingly, rapid evolution was evident in the RIPK3 RHIM domain but not detectable in homologous domains across primate *TRIF*, *DAI,* and *RIPK1* (Fig 3D). Indeed, *RIPK3* showed elevated rates of substitution by PAML analysis at two sites immediately adjacent to the determined RHIM EspL cleavage motif – G<u>DN</u>/NYL where “/” indicates cleavage; the sites 462D and 463N are rapidly evolving (Fig 3D). The evolutionary signatures of necroptosis RHIM domains may indicate modes of RIPK3 activation and/or functions independent of the RIPK1, TRIF, and DAI RHIM domains.

### Poxviral antagonism of the RIPK3:MLKL interface serves as a model for antagonistic coevolution

The rapid evolution signature of RIPK3 near the hydrophobic pocket indicates antagonism, likely by pseudosubstrates (Fig 3C). Indeed, a viral copy of MLKL (vMLKL), which is encoded by numerous poxviruses has been recently identified [31]. Yet, the origins and evolution of these vMLKLs remain unknown. We identified twenty-seven distinct copies of vMLKL in the sequence database across the genomes of *Avipoxvirus*, Clade II, along with two other poxvirus species that infect bats [eptesipoxvirus (EPTV) and hypsugopoxvirus (HYPV)] (Fig 4D). vMLKL lacks the N-terminal bundle and brace (NBB) domain but maintains the C-terminal kinase-like domain, to which RIPK3 binds to phosphorylate the host cellular homolog [31] (Fig 4A). Ectopic expression of two vMLKLs from poxvirus species that primarily infect mice were demonstrated to competitively inhibit both mouse and human RIPK3 in binding and cell-culture necroptosis assays [31]. Superposition of a homology model of myxoma virus (MYXV) MLKL onto published crystal structures for mouse MLKL [49] and human MLKL [50] indicated striking structural overlap with human and mouse MLKL (Fig 4B). This occurs despite a broad-range of amino acid divergence between these host and viral MLKLs (16-25% amino acid identity of the virus homologs to mouse and human MLKLs).

**Fig 4.**
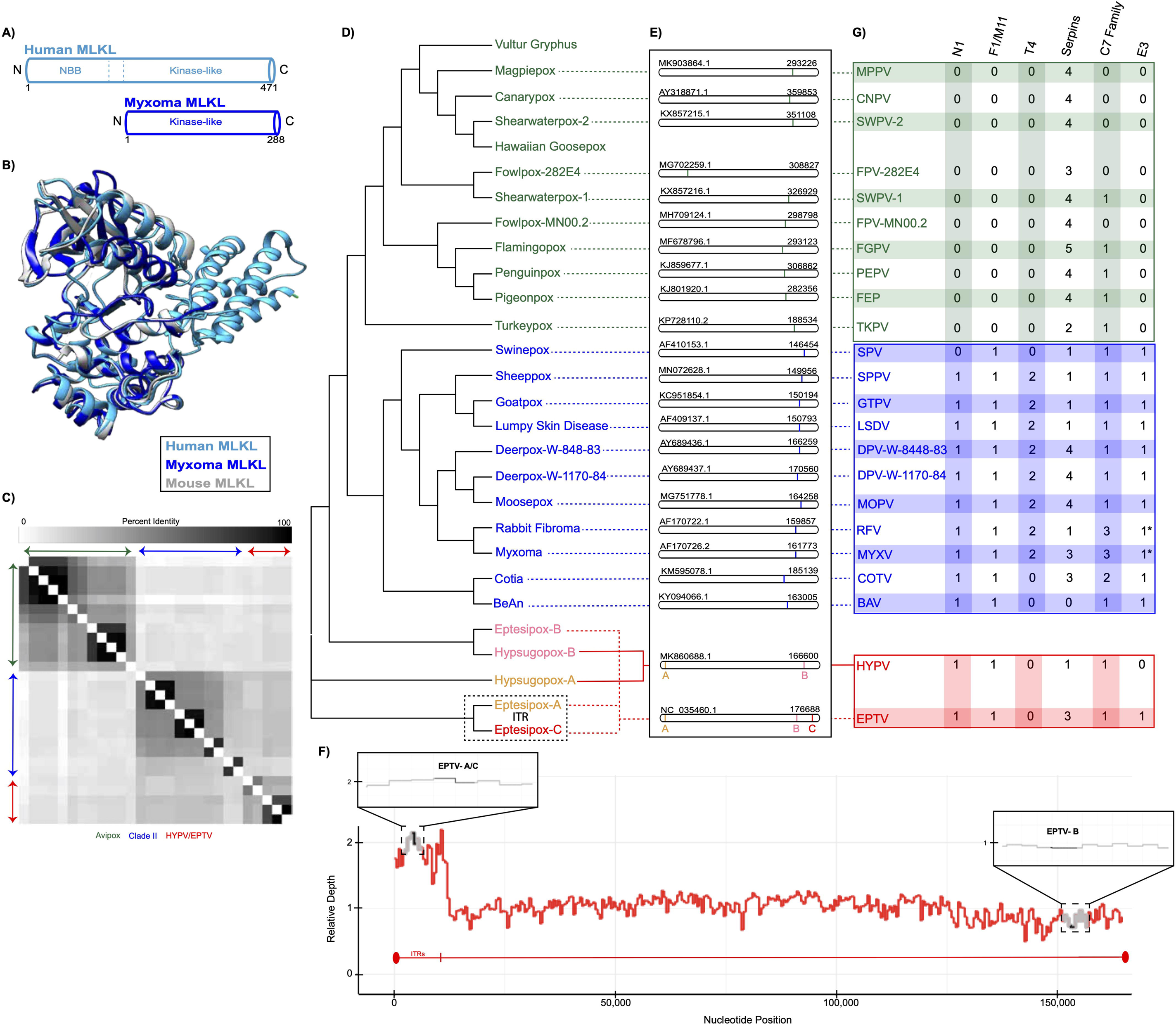
vMLKL mimics are encoded in *Avipoxvirus*, Clade II, eptesipoxvirus, and hypsugopoxvirus genomes. **A)** Protein structures of human MLKL (light blue) and myxoma v(irus)MLKL (dark blue). **B)** Homology model for myxoma vMLKL (dark blue) predicted using Swiss-Model and aligned onto previously published crystal structures for hMLKL (light blue) and mMLKL (silver). **C)** Heatmap comparing percent identities of vMLKL copies across poxvirus species which were included in this study. **D)** Cladogram depicting distinct species containing a vMLKL copy with *Avipoxvirus* (green), Clade II (blue) and other poxvirus species (yellow, pink and red) separated according to genus. Note that eptesipoxvirus and hypsugopoxvirus contain multiple copies of vMLKL and are labeled based on their position in the chromosome – A (yellow), B (pink), C (red). Eptesipoxvirus (black dashed box) contains vMLKL copies located on the inverted terminal repeats (ITR) represented by A (yellow) and C (red). **E)** Location of individual and multiple copies of vMLKL across poxvirus genomes. Multiple copies of vMLKL are highlighted (yellow, pink, and red) in eptesipoxvirus and hypsugopoxvirus. Chromosomal cartoons created in CoGeBlast represent scaled locations of the vMLKL copies for each of the poxvirus species. The genome accession numbers are located on the top left corner of the chromosomes, and the nucleotide lengths are located on the top right corner of the chromosomes. **F)** Multiple copies of vMLKL in the eptesipoxvirus genome are supported by relative depth analysis. The distribution of reads that align with the ITR and non-ITR regions is shown. Zoomed in images, represented by black boxes, show the relative depth of the EPTV vMLKLs (black) and the two flanking upstream and downstream genes (gray). The ITR relative depth mean is 1.852 and the median is 1.899. The non-ITR relative depth mean is 1.000 and the median is 1.025. **G)** Repertoire of poxvirus apoptosis and necroptosis antagonists explained, in part, by evolutionary distribution of vMLKL. Only proteins predicted to be functional are shown. Two exceptions, found in myxoma virus and rabbit fibroma virus, are represented by an ‘*’ and have N-terminally truncated E3 copies. In depth analysis can be found in S2 File.

In agreement with the strong structural overlap, characterization of vMLKL sequences indicate conservation of functional residues for human and mouse MLKL including ATP binding, hydrogen-bonding, and protein stability (S2A Fig) [19-21, 49-51]. Several amino acid sites displaying positive selection for MLKL (Figs 2 and 3) overlapped or were in close proximity to sites for hydrogen bonding and protein stability, as demonstrated by the amino acid alignment (S2A Fig). Of note, there are several intriguing examples of specific residue conservation between the primate positively selected sites and the viral copies. One example is human MLKL F386 (F373 mouse), which has been shown to be necessary for RIPK3/MLKL complex formation [31]. This position exclusively has tyrosine, phenylalanine, or histidine amino acid variants across the sampled primates, with this pattern of aromatic residues evident and conserved to Aves MLKL virologs (S2B Fig). Consistently, poxvirus virologs have also maintained only aromatic (phenylalanine or tyrosine) amino acids at the homologous position (S2A Fig). These data suggest a key role for site 386 in host defense but also in counteraction by viral mimics [52].

To further understand the evolutionary relationships between poxvirus vMLKLs, we performed detailed sequence and phylogenetic analyses across corresponding viral genomes. Full-length vMLKLs (excluding partial Vulture gryphus poxvirus and Hawaiian goosepox virus sequences) display 18%-100% amino acid identity relative to each other and cluster in a manner resembling known poxviral species relationships (Fig 4C). Unexpectedly, this analysis identified two distinct vMLKL copies (e.g. paralogs), which differ in amino acid identity, in the genomes of both eptesipoxvirus (EPTV vMLKL-A vs. EPTV vMLKL-B : 45% amino acid identity) and hypsugopoxvirus (HYPV vMLKL-A vs. HYPV vMLKL-B : 43% amino acid identity). Analysis of the location of the different vMLKLs in each genome was conducted using CoGeBlast, which allows for individual hit visualization. The results suggested distinct genomic locations for these genes (Fig 4E). Specifically, a majority of vMLKLs are located primarily on the right arm of the poxvirus genome with EPTV having two copies that map to two distinct loci in this region. In addition, both HYPV and EPTV have one additional copy that maps to the left arm of the poxvirus genome. Interestingly, two of the vMLKLs in EPTV are present on the inverted terminal repeat (ITR), potentially representing more recent acquisitions. Likewise, fowlpox virus-282E4 vMLKL appears to be at a distinct inverted locus relative to other *Avipoxvirus* species on the left arm of the genome (Fig 4E).

To confirm additional vMLKL copy numbers present in the eptesipoxvirus genome, we next performed relative depth analysis, with approximately 1000 - fold genome coverage (Fig 4F, S3 Fig), using the raw reads from the genome sequencing. By mapping reads across the unique genome and one reference ITR, we found that EPTV – A/C has a relative depth of two and EPTV – B has a relative depth of one, which supports the presence of three copies in the eptesipoxvirus genome (Fig 4F). As a control, we also conducted a similar analysis using reads from a recent myxoma virus sequencing project [53] and found that the vMLKL relative depth remained near one, indicating the presence of only one genomic copy (S3 Fig). These data demonstrate that vMLKLs have different evolutionary histories and are likely not orthologous.

### Phylogenetic distribution of vMLKLs can account for variable conservation of E3 across species

Poxviruses encode numerous immunomodulators to subdue and circumvent host defenses including apoptosis. As a majority of poxviruses encode several characterized anti-apoptotic proteins [54, 55] (Fig 4G, S4 Fig, S2 File) including the serpin class of caspase pseudosubstrates [56, 57], these viruses would be expected to trigger necroptosis. Indeed, the model poxvirus vaccinia has been shown recently to trigger necroptosis through activation of DAI/ZBP1 in cultured cells and to counteract this response using the Z - nucleic acid binding (Zα) domain of the E3 protein [32]. E3 is considered a key host-range gene that antagonizes several host defenses. Interestingly, some poxviruses that encode a repertoire of anti-apoptotic proteins lack E3 or possess truncated E3 ORFs. For example, the *Avipoxvirus* family does not possess any orthologs of E3 (Fig 4G, S2 File) and several Clade II poxviruses, including the well- characterized myxoma virus, are also predicted to have N-terminally truncated E3 copies which lack the Zα domain that is important for antagonism of DAI [55] (Fig 4G, S2 File). To gain insights into how a subset of poxviruses may circumvent necroptosis given the presumed ability to trigger apoptosis, we compared the breadth of known anti- apoptotic proteins, including serpins, across the poxvirus phylogeny relative to vMLKL (Fig 4G). As vMLKLs are present in *Avipoxvirus*, hypsugopoxvirus, myxoma virus and rabbit fibroma virus – the maintenance of this pseudosubstrate may account for the tolerance of these species to a missing or truncated E3 protein.

### Repeated gains and losses of vMLKL over poxvirus evolution

Our data illustrates that multiple vMLKL copies may exist in a given poxvirus genome and that not all of the copies across the genomes may be orthologous. To determine the relationship of poxvirus vMLKLs, we performed a synteny analysis using five flanking upstream and downstream genes across the entire phylogeny (Fig 5, S3 File). These data indicate that while each vMLKL within *Avipoxvirus* and Clade II are flanked largely by the same upstream and downstream genes, vMLKLs between the species are flanked by different genes. The Clade II and *Avipoxvirus* species have distinct copies as indicated by different upstream and downstream genes. Furthermore, the lack of vMLKL in Yatapoxvirus species (Yaba monkey tumor virus and Yaba-like disease virus), which belong to Clade II, revealed that this loss is likely due to a deletion event consisting of multiple upstream and downstream genes in these lineages (Fig 5, S3 File).

**Fig 5.**
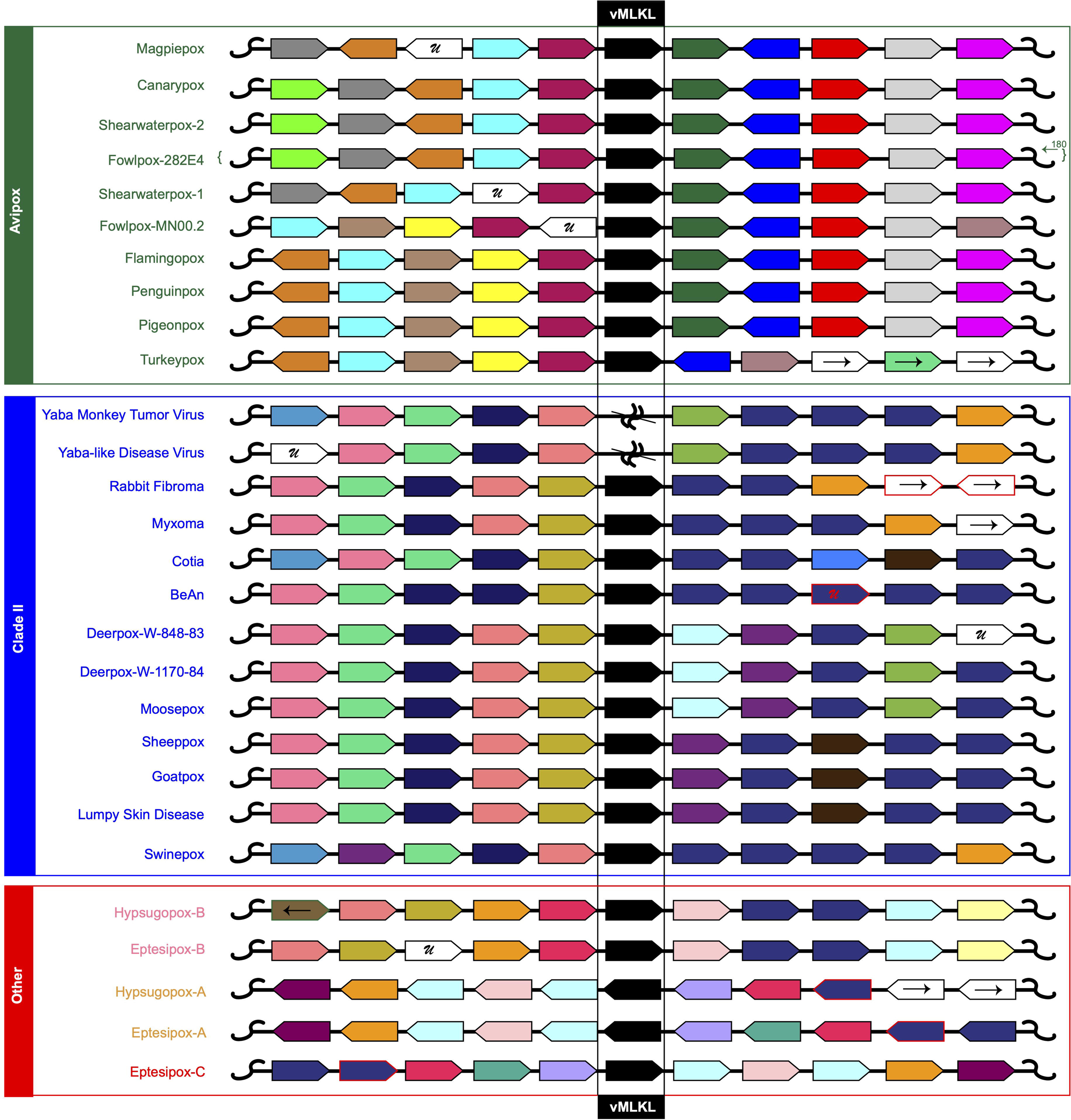
High resolution synteny analysis suggests diverse evolutionary histories for vMLKL. Analysis of upstream and downstream flanking genes indicate vMLKLs (black) reside at different locations across poxvirus genomes. Protein homology was determined using reciprocal BLAST hits for the 5 genes upstream (left of vMLKL) and 5 genes downstream (right of vMLKL) of vMLKL. While a gene window of 11 is shown here, an expanded analysis is found in the S3 File. Genes oriented to the right represent 5’-3’ directionality while genes oriented to the left represent 3’-5’ directionality. The letter “U” represents a gene that is either unique to the individual species or unique to the group of poxviruses that have maintained a vMLKL copy. Gene blocks that have homologous genes (colored borders) that are outside the scope of this figure have arrows pointing to the left (representing upstream) or to the right (representing downstream). Two Yatapoxvirus species (Yaba monkey tumor virus and Yaba-like disease virus) were included as they belong to clade II but lack vMLKL. 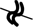 represents a deletion event. Species defined as “Other” (red) are either unclassified (hypsugopoxvirus) or part of vespertilionpoxviruses (eptesipoxvirus).

This analysis also provided insights regarding the multiple vMLKL copies in eptesipoxvirus and hypsugopoxvirus. First, this analysis indicated that EPTV-B (EPTV- WA-166) and HYPV-B (QDJ95132.1) are orthologous to each other and are likely orthologous to the Clade II copies, as evidenced by some overlap in syntenic gene neighbors (Fig 5). Second, EPTV-A (EPTV-WA-006) and HYPV-A (QDJ94987.1) share gene neighbors indicating that these sequences are likely orthologs. EPTV-A shares 100% identity with EPTV-C (EPTV-WA-186) with both annotated as being located in the inverted terminal repeats (ITRs). Consistently, EPTV-C has the same gene neighbors as EPTV-A albeit inverted. The copies are labeled corresponding to their location starting from the left arm of the poxvirus genome – e.g., A, B, C.

To further assess the evolutionary relatedness of vMLKLs, we performed whole genome comparisons using CoreGenes (https://coregenes.ngrok.io/). This data was visualized with RIdeogram [58] and focused on the copies encoded by *Avipoxvirus*, Clade II and the two poxvirus species that primarily infect bats. Indeed, we found that canarypox virus vMLKL and myxoma virus vMLKL localize to distinct chromosomal locations with no shared surrounding regions of synteny (S5A Fig). In agreement with our local synteny analysis above, the locus encoding Clade II vMLKL was likely lost in a Yatapoxvirus ancestor following divergence as evidenced by global as well as local synteny surrounding the vMLKL locus (S5B Fig). Lastly, we compared eptesipoxvirus with myxoma virus (S5C Fig). The upstream regions of synteny of vMLKL indicate that MYXV vMLKL is orthologous to the EPTV-B vMLKL copy, which is further supported by MYXV vMLKL not being in the ITRs corresponding to EPTV-A/C. These data suggest vMLKLs are at distinct loci in poxvirus genomes as a result of repeated gains and losses during poxvirus genome evolution.

### Phylogenetic analysis reveals origination of vMLKL copy from one horizontal gene transfer event

The distinct copies of vMLKLs may have originated due to either 1) a single transfer event from a host genome to an ancestral poxvirus followed by diversification or multiple independent transfer events from presumably distinct host genomes to different poxviruses. To address this, we performed a phylogenetic analysis using amino acid sequences of vMLKLs with thirty-eight diverse host MLKLs. If distinct vMLKLs were acquired via independent events, placement of viral sequences would vary across the tree in a manner where they pair with different host sequences. Such topology could reflect the source genome or that of a related, perhaps ancestral, animal host. In contrast, if vMLKLs originated from a single transfer event, the tree topology would group these sequences into one clade. Phylogenetic analysis using PhyML [59] revealed that vMLKL virologs cluster as one group distinct from animal vMLKLs (Fig 6). These data support a single transfer event from an ancestral host genome as the most parsimonious origin for vMLKL (Fig 6, S4 File). The topology of the poxvirus branches mirrors that of known species relationships. For example, all of the *Avipoxvirus* species cluster together. Likewise, the Clade II species cluster together broadly, as well as at a finer level such as in the *Leporipoxviruses* (e.g. MYXV and RFV). The tree topology also supports the synteny analysis with HYPV-B and EPTV-B being most closely related along with EPTV-A/C and HYPV-A vMLKL clustering distinctly.

**Fig 6.**
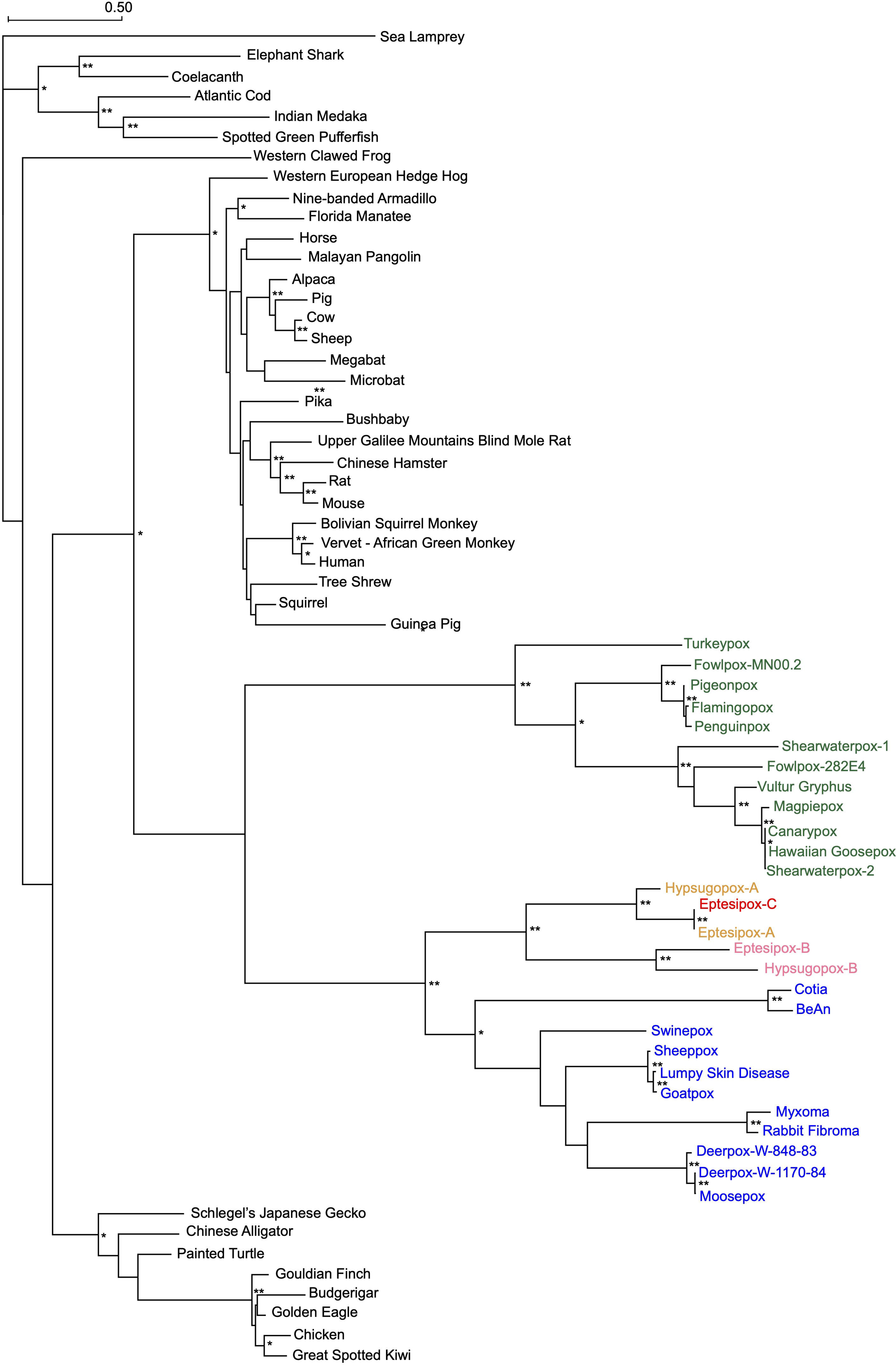
Phylogenetic analysis suggests a singular origin for non-orthologous poxvirus MLKLs. A maximum-likelihood phylogenetic tree of diverse host and poxviral MLKL amino acid sequences produced using PhyML [59] is displayed. Consistent with a single horizontal gene transfer event, *Avipoxvirus* (green), Clade II (blue), and other poxvirus sequences (yellow, pink, and red) cluster together. Branch lengths represent amino acid changes per site. 100 bootstrap replicates were performed with branch support > 70% (*) or 90% indicated (**). The alignment of MLKL sequences was performed using MAFFT and can be found in the S4 File.

Still, it remains unclear whether Clade II vMLKLs represent a distinct or orthologous locus to Avipox vMLKL and which vMLKL locus represents the ancestral locus. Indeed, our data for Clade II vMLKL could be explained, in part, by either 1) a duplication of the Avipox vMLKL locus followed by a deletion event of the original locus in the Clade II ancestor or 2) that Avipox and Clade II vMLKL are orthologous but divergence no longer allows for syntenic comparison between these species. We assessed the likelihood of these evolutionary histories by analyzing patterns of amino acid variation shared between vMLKLs and mammalian MLKLs, which cluster closest to vMLKLs (Fig 6). We evaluated innovations specific to vMLKLs but not host MLKLs across our alignment. We identified seventeen amino acid residues largely conserved between Avipox vMLKLs and mammalian MLKLs (Fig 7) but not Clade II, EPTV and HYPV vMLKLs. Conversely, different variants at the same sites are shared across Clade II, EPTV and HYPV vMLKLs but not Avipox or mammalian MLKLs. These data support the hypothesis that Avipox vMLKL is the original locus and that Clade II vMLKL is a distinct copy but derived from the Avipox vMLKL presumably through duplication. Collectively, these data illustrate a volatile history of vMLKL since its acquisition by an ancestral *Avipoxvirus*.

**Fig 7.**
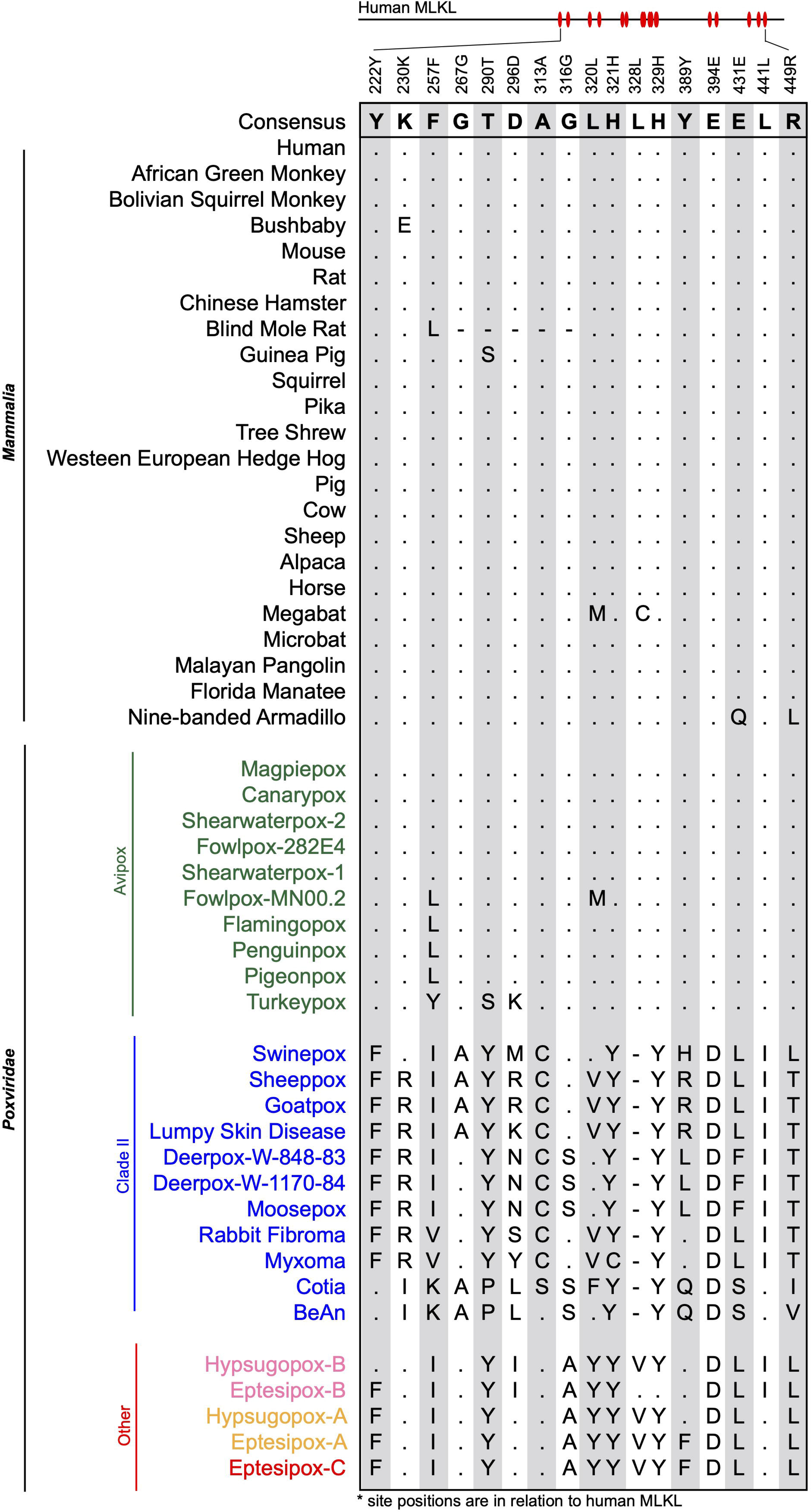
Amino acid variant analysis suggests Avipox vMLKL is the original locus, and Clade II vMLKL is an independent, but derived copy. A set of informative amino acid positions were identified in the alignment of mammalian and vMLKL sequences. Residues identical to consensus are shown as “.” while “-” represents a gap in the sequence alignment. Amino acid site positions (shown above the alignment) are in relation to human MLKL. Alignment can be found in the S4 File.

## Discussion

Studies of host-pathogen co-evolution can reveal new insights that forge infection outcomes. These include vulnerabilities in host defense and determinants that shape pathogen host range as well as zoonoses. In addition, analysis of host and pathogen effectors can illuminate new evolutionary paradigms such as the evolutionary trajectories of viral mimics. Here, we focused on the emerging regulated cell death program of necroptosis that is largely presumed to be a “back-up” response. Our data, particularly the robust signatures of positive selection in RIPK3/MLKL, mirrors that of key immune factors like cGAS, PKR, and OAS1 [37, 38]. These features support the essentiality of RIPK3/MLKL in immunity during primate evolution. In addition, the maintenance, as well as gains and losses of vMLKL throughout poxvirus evolution, showcases the need to actively counteract the necroptosis program.

### Proposed model for vMLKL acquisition

The findings from our evolutionary analysis for vMLKL are consistent with a model consisting of a series of gains and losses of this virolog over time (Fig 8A). Our data paired with those of Petrie *et al*., which also suggested a host origin for vMLKL [31], support the notion that this gene was initially acquired via one horizontal gene transfer event (Fig 6). As vMLKL mirrors a truncated copy of cellular MLKL lacking introns, we hypothesize that this transfer likely occurred via an RNA intermediate. The N-terminal truncation in vMLKL could have transpired either during or after integration. This copy at the original site of integration has been maintained through evolutionary time in *Avipoxvirus* species and represents locus 1. Following the Avipox ancestral divergence from a common ancestor shared with Clade II and HYPV/EPTV species, locus 1 appears to have been duplicated to produce locus 2. We hypothesize that locus 1 was deleted while locus 2 was maintained in the shared ancestor Clade II/EPTV/HYPV, and is present today in most contemporary genomes of Clade II poxviruses and EPTV-B/HYPV-B. Yet, it remains possible - due to extensive rearrangements resulting in the degradation of any detectable synteny by either method we employed here - that locus 1 across Avipox represents the Clade II/EPTV/HYPV vMLKL. In Yatapoxviruses, locus 2 vMLKL was lost in what appears to be a multi-gene deletion event (Fig 5, S5 Fig). Following divergence of the EPTV/HYPV ancestor from Clade II poxviruses, locus 2 was duplicated to a region in the ITR with a subsequent duplication occurring in the other ITR. Both vMLKL copies in the eptesipox ITRs are supported by our relative depth analysis (Fig 4F, S3 Fig). A similar analysis was attempted but not possible with the hypsugopox genome data. Thus, vMLKL resides at four distinct loci in poxvirus genomes across the phylogeny.

**Fig 8.**
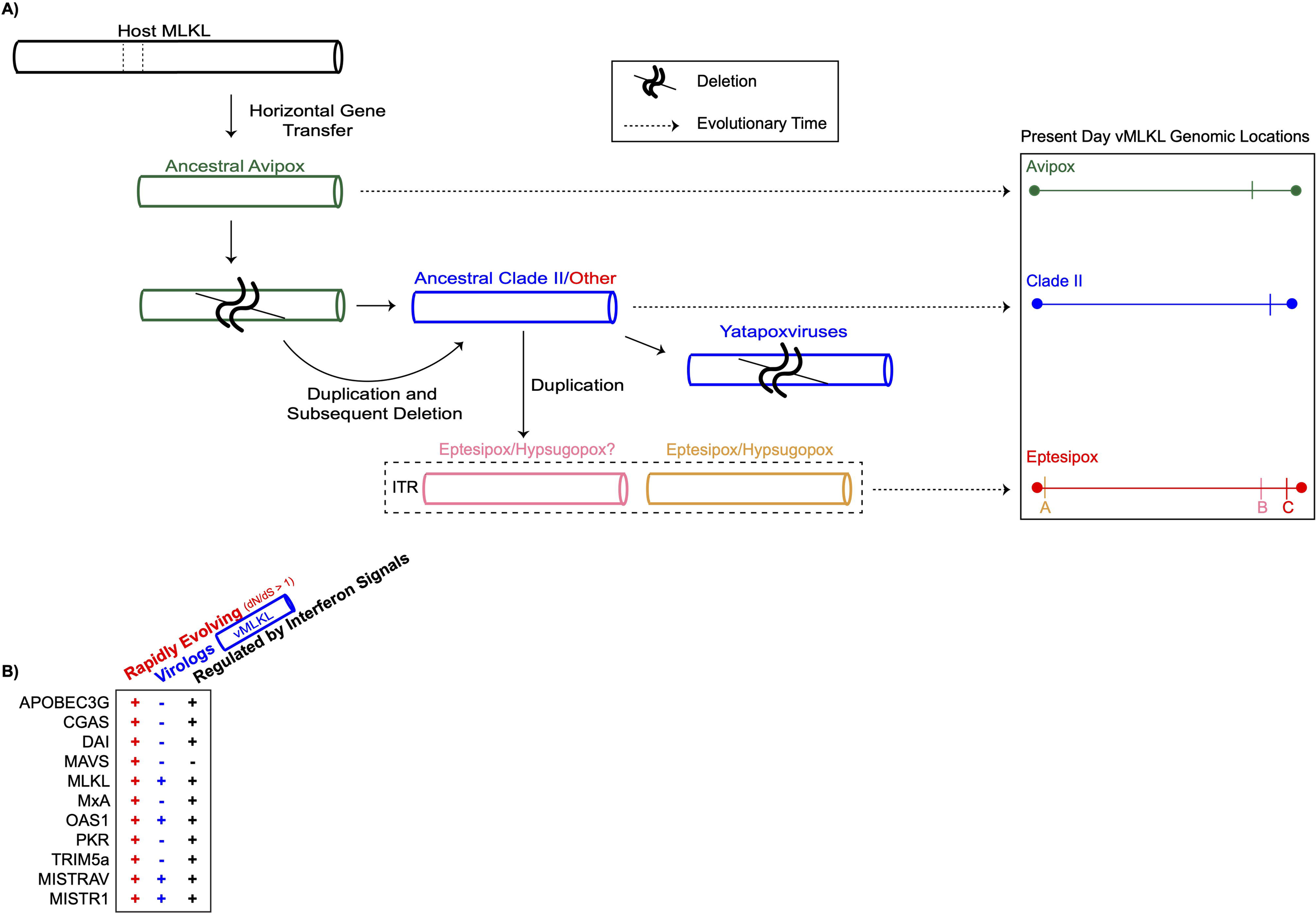
Volatile evolution of MLKL and hallmarks associated with key host defense factors indicate that necroptosis is a pivotal, not auxiliary, immune defense. **A)** Model of poxvirus MLKL acquisition and evolution across lineages. Data indicate that vMLKL was likely acquired by a vertebrate host presumably through a single horizontal gene transfer event to a poxvirus ancestor (Fig.6). This copy at the original site of integration, located on the right arm, has been maintained in *Avipoxvirus* (green) genomes over time to present day (Fig 4). A duplication followed by a deletion event occurred in the common ancestor of Clade II (dark blue) and species classified as “Other” (red) following divergence from *Avipox* (Figs 5 and 7, S5 Fig). This second copy also on the right arm has been maintained until the present in all Clade II species but lost in Yatapoxvirus lineages (Fig.5). Subsequently, the second copy was duplicated to the ITR in the ancestor of eptesipoxvirus and hypsugopoxvirus, which is maintained in present-day lineages. While hypsugopoxvirus ITR sequence was not included in the final genome assembly, our synteny analysis (Fig 5) suggests vMLKL is also located on an ITR (dashed box) in this lineage. “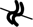” represents predicted deletion events. Dashed arrows represent the progression of evolutionary time (left to right). Scaled chromosomal locations of the present day vMLKL copies are represented on the right. **B)** MLKL displays all three key hallmarks often common to key host defense factors but unique to OAS1 [38, 39, 61], MISTRAV, and MISTR1 [62]. These factors are rapidly evolving, possess a viral homolog/virolog, and are regulated by immune signals (includes upregulation and downregulation).

### Positive selection signatures in the necroptotic signaling pathway are localized to the RIPK3/MLKL axis

The signatures of rapid evolution in RIPK3 and MLKL resemble “molecular arms races” dynamics with pathogen-encoded inhibitors [48] (Fig 2). The broad distribution of rapidly evolving sites in both RIPK3 and MLKL is indicative of multiple, undefined antagonists that have existed over evolutionary time (Fig 3). Interestingly, our analysis across the core circuitry of necroptosis enabled two key observations. First, strong selective pressure has been primarily exerted on the downstream components of the pathway where upstream signaling converges (Figs 2 and 3). Second, our evolutionary analysis across the pathway allowed the identification of rampant positive selection exclusive to the RHIM-domain of RIPK3 but not the RHIM-domains of the upstream activators DAI, TRIF, or RIPK1 (Fig 3). This is extremely striking given the elevated dN/dS ratios for RIPK3, but not the other factors, at amino acid positions that define a demonstrated cleavage site (Fig 3D) for the conserved bacterial protease EspL which is capable of cleaving RHIM domains across this pathway. Notably, rapid evolution in the adaptor protein MAVS has been shown to shape the ability of the hepatitis protease NS3/4A to cleave this target [60]. Our data suggest that there may be RIPK3 activity that is independent of activation by RHIM homotypic interactions (e.g., DAI/TRIF/RIPK1), which is counteracted by EspL-like proteases. One possible explanation for this signature is the presence of a yet undefined upstream activator of RIPK3, which lacks a RHIM-domain, that is triggered by pathogenic bacteria. Given the data, we are unable to infer whether this predicted activity of RIPK3 would also activate MLKL. Nevertheless, a pathway perspective combined with evolutionary insights can drive new avenues of investigation.

### RIPK3/MLKL as a model for the maintenance of compatible, functional interactions

Our study describes volatile evolution of vMLKL in poxvirus genomes. This work extends on the recent discovery of this mimic, which showed two vMLKLs are capable of counteracting necroptosis in cell culture [31]. Our integrated comparison of established poxvirus immunomodulators with the phylogenetic distribution of vMLKL appears to resolve the long-standing conundrum of how some poxvirus species persist with N-terminally truncated or completely absent E3 ORFs (Fig 4F). In addition, the lack of vMLKLs in orthopoxviruses can be explained, in part, by the recent identification of vIRD encoded by this genus, which is an ankyrin-repeat domain containing protein that targets RIPK3 for degradation [35]. The resolution of the evolutionary history of vMLKL in our study may serve as a resource in guiding the functional analysis of host and viral MLKL variants and their impact on RIPK3 binding and activity. It has not escaped our attention that the RIPK3/MLKL axis, in part, resembles another well-defined host defense axis - PKR/eIF2 α - which also consists of a kinase and its substrate counteracted by a different poxvirus pseudosubstrate (K3L) [37]. However, a major difference between the two systems is that RIPK3 and MLKL are rapidly evolving, in contrast to PKR rapidly evolving and eIF2 α being ultraconserved to yeast. This is interesting given the PKR/eIF2 α system has been useful in understanding how a rapidly evolving factor maintains its interaction with a conserved factor while evolving to escape antagonism [37]. Perhaps RIPK3/MLKL may be powerful in understanding how two key host defense factors maintain their essential interaction to drive necroptosis while escaping pathogen-mediated inhibition, particularly pseudosubstrates similar to vMLKL.

### MLKL is a pivotal immune factor

Our study defines MLKL, which functions as the executioner of necroptosis, as a factor that 1) is (up)regulated by immune signals (Fig 2B) [43–45], 2) has a viral homolog/virolog (Figs 4,5,6 and 7)[31] and 3) is rapidly evolving (Figs 2 and 3). While several key host defense factors typically display two of these signatures, to our knowledge, only a select few are published which display all three of these hallmarks (Fig 8B): OAS1 [38, 39, 61], and the recently described MISTRAV and MISTR1 [62]. OAS1 is one of the first interferon-stimulated genes identified and functions in host defense against numerous, diverse viruses [63]. MISTR proteins appear to interface with electron transport chain complexes to regulate stress responses and cellular adaptation during infection [62]. We hypothesize this collection of three hallmarks may serve as a guide to identify other pivotal host defense factors. Collectively, we interpret this rare combination of signatures that are associated with OAS1 and now MLKL as strong evidence that necroptosis is an indispensable, not auxiliary, host response that has shaped infection outcomes over millions of years.

## Methods

### Protein modeling

Published crystal structures of m(ouse) RIPK3 (PDB: 4M66), mMLKL (PBD:4M68) and co-crystal structure for mRIPK3:mMLKL (PDB: 4M69) were used for modeling following pairwise alignment with h(uman)MLKL (PDB: 4BTF) or the predicted structure of hRIPK3 [20, 49]. Swiss-Model was used to predict the structure of hRIPK3 and myxoma v(iral)MLKL [64]. UCSF Chimera (https://www.cgl.ucsf.edu/chimera/) was used to perform analysis and visualize structures including rapidly evolving sites [65].

### Positive selection analysis

Nucleotide sequences were obtained from NCBI and Ensembl databases (S1 File). Multiple sequence alignments (MSA) were performed using MAFFT iterative refinement method FFT-NS-i and visualized using Geneious Prime 2020.1.2 (https://www.geneious.com/) [66]. Indels were removed through manual trimming. dN/dS lineage estimates were obtained using sample primate gene MSA and phylogenetic tree [created using previously described primate lineage relationships [67]]. These served as input for FreeRatio analysis implemented in PAML [42]. Nucleotide substitution (NS) site analysis was performed with PAML using both F3×4 and F61 codon frequency models. MEME and FUBAR analysis were performed using Datamonkey (https://www.datamonkey.org/) to predict positively selected amino acid sites [46, 47, 68]. The S1 File contains additional findings from this analysis.

### Identification of poxvirus vMLKL

Virologs to human MLKL were identified using tBLASTn and/or BLASTp [69]. Sequences were selected based on returned BLAST parameters (total score, E value and percent identity) and the phylogenetic clustering of the sequences using PhyML [59]. Scaled individual hit visualization was constructed using CoGeBlast [70].

### Relative depth analysis

Eptesipox raw sequencing reads, published in Tu *et al.,* were obtained directly from Dr. Chris Upton (University of Victoria, Victoria BC, Canada) and Dr. Yoshinori Nakazawa (CDC, Atlanta, Georgia) [71]. NC_035460.1 was used as the eptesipoxvirus reference sequence to map the reads. Myxoma virus raw sequencing reads were obtained from European Nucleotide Archive (https://www.ebi.ac.uk/ena/browser/home), from Project PRJNA513218 [53]. Accession IDs used for this analysis are SRR8402032, SRR8402033, SRR8402034, and SRR8402035. As a control, AF170726.2 was used as the myxoma virus reference sequence to map the reads. Raw sequencing reads were processed using Trimmomatic-0.39 [72] to remove adaptors. Tanoti short read aligner (https://bioinformatics.cvr.ac.uk/software/tanoti/) was then used to map reads to the reference genome. SAMtools [73] was used to convert between file formats, merge files and calculate coverage per basepair. Bins of 500 nucleotides were created for relative depth analysis, which was calculated by dividing the bins by the mean of the non-ITR genomic region. Data was visualized using gplots [74].

### Phylogenetic analysis

Multiple Sequence Alignments were performed using MAFFT, with iterative refinement method L-INS-i, on amino acid sequences retrieved from NCBI and Ensembl databases (S4 File) [66]. Maximum Likelihood trees were inferred by PhyML using 100 bootstrap values [59].

### Percent amino acid identity heatmap

Multiple Sequence Alignments were performed using MAFFT, with iterative refinement method L-INS-i, on amino acid sequences retrieved from NCBI database (S2 File) [66]. The R package gplots was used to create the heatmap found in Fig 4C [74, 75].

### Poxvirus RCD-associated host range gene identification

Host range genes were identified using BLASTp and tBLASTn. Sequences were confirmed using reciprocal BLAST hits. Protein families were found using genomic database annotations and published results for included poxvirus genomes. Domain classification for homologs and protein families were confirmed using NCBI conserved genome database [76]. Results were compared to previous homolog studies performed by Bratke *et al*. 2013 and Haller *et al.* 2014, and can be found in S2 File [54, 55]. Proteins truncated at the N- and C-termini were not included in Fig 4G but can be found in the S2 File with appropriate label. An exception is necroptosis inhibitor E3, which is predicted to be N-terminally truncated in *Leporipoxviruses* [55]. Genes residing in the inverted terminal repeats (ITR) are counted as separate genes and thus two instances in order to stay consistent with the Fig 4C initial analysis, where the eptesipoxvirus ITR is included and analyzed. The p28/N1R and Ankyrin-repeat families were annotated using NCBI database annotations. The presence of these domains were confirmed using NCBI conserved genome database or via BLASTp [76]. This analysis includes all proteins possessing a p28/N1R-like or Ankyrin-repeat domain and does not predict if these proteins are functional and/or likely truncated.

### High resolution synteny analysis

Reciprocal BLAST hits (RBH) were performed for coding sequences upstream and downstream of vMLKL genomic locations across poxvirus species included in this study. Results can be found in the S2 File. Genomic database annotations and the NCBI conserved genome database were used to confirm results [76]. Gene blocks representing Ankyrin-repeat containing proteins without one distinct hit, but conserved homology across poxvirus species are colored navy.

### Whole genome comparison

Coregenes 5.0 (https://coregenes.ngrok.io/) was used to compare selected poxvirus genomes [77]. Matches were identified using bidirectional best hit with an e-value = 1×10^5^. RIdeogram [58] was used to visualize the output obtained from Coregenes 5.0.

### Cell lines

HT29 and HeLa cell lines were obtained from ATCC. Both cell lines were cultured in Corning DMEM with L-Glutamine, 4.5 g/L Glucose, and Sodium Pyruvate supplemented with 10% FBS and 1X Gibco Antibiotic-Antimycotic solution. Cell lines were maintained at 37°C in a humidified incubator at 5% CO_2_.

### Cell culture treatments

Cells were treated with either human IFNα [1000 U/mL (PBL Assay Science, USA) or human IFNγ [1000 U/mL (ThermoFisher, USA)] diluted in DMEM and incubated for 24 hours. After 24 hours, cell lysates were harvested for western blots as described below.

### Western blot analysis

Cells were collected using RIPA Lysis and Extraction Buffer (ThermoFisher, 89901) supplemented with Protease and Phosphatase Inhibitor Cocktail (Abcam, ab201119). Protein concentrations were measured using a Bradford assay. For all blots, 5 _µ_g of protein lysates were loaded onto an SDS-PAGE gel and wet-transferred to 0.45 _µ_M Immobilon-P PVDF membrane (Millipore, IPVH00010) at 200 mA for 90 minutes. Membranes were blocked with blocking buffer (5% milk in TBST) for 30 minutes at RT. Membranes were incubated in primary antibodies at 4°C overnight. The following primary antibodies used: MLKL (GeneTex, GTX107538), STAT1 D1K9Y Rabbit mAb (Cell Signaling Technology, 14994S), P-STAT1 Tyr701 58D6 Rabbit mAb (Cell Signaling Technology, 9167S) and β Actin (Abcam, ab49900). Membranes were washed three times with TBST for 5 minutes and incubated with Goat Anti-Rabbit IgG (Bio-Rad, 170-6515) for 1 hour at RT. Membranes were washed three times with TBST and incubated with Clarity Max Western ECL Substrate (Bio-Rad, 1705062). Following incubation, blots were imaged using the Chemidoc MP Imager (Bio-Rad).

## Acknowledgments

We express gratitude to other members of the Hancks Lab for feedback and discussion through the course of this project. We also thank Dr. Anant Gharpure and Dr. Sherry Haller for their comments on the manuscript. We also thank Dr. Chris Upton and Dr. Yoshinori Nakazawa as well as Dr. Chiara Chiapponi and Dr. Davide Lelli for their assistance with the eptesipoxvirus and hypsugopoxvirus data, respectively.

## Supporting information

**S1 Fig.**
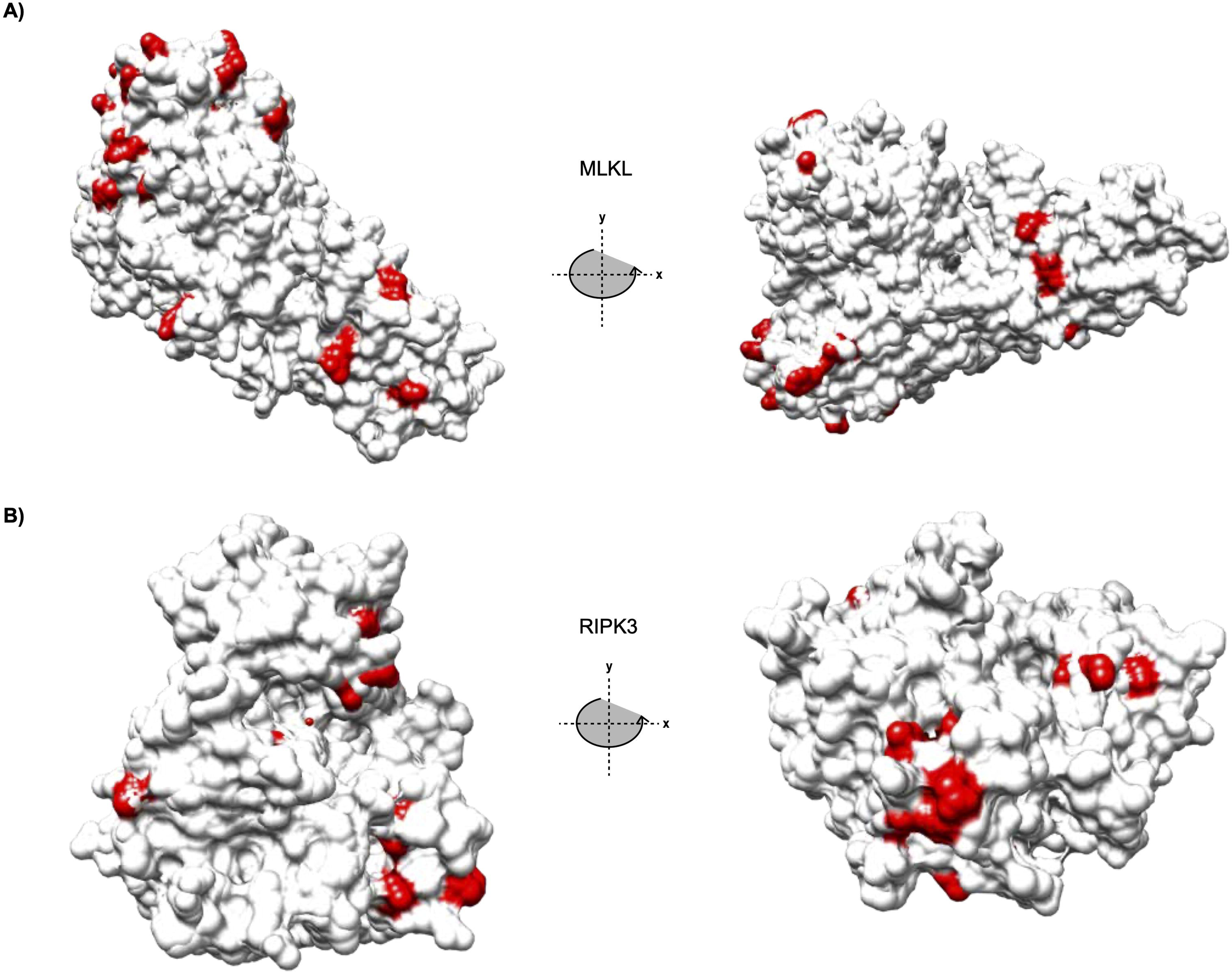
MLKL and RIPK3 rapidly evolving sites localize to protein surfaces. Positively selected sites (red) identified in Fig 3 mapped onto filled crystal structures of RIPK3 and MLKL.

**S2 Fig.**
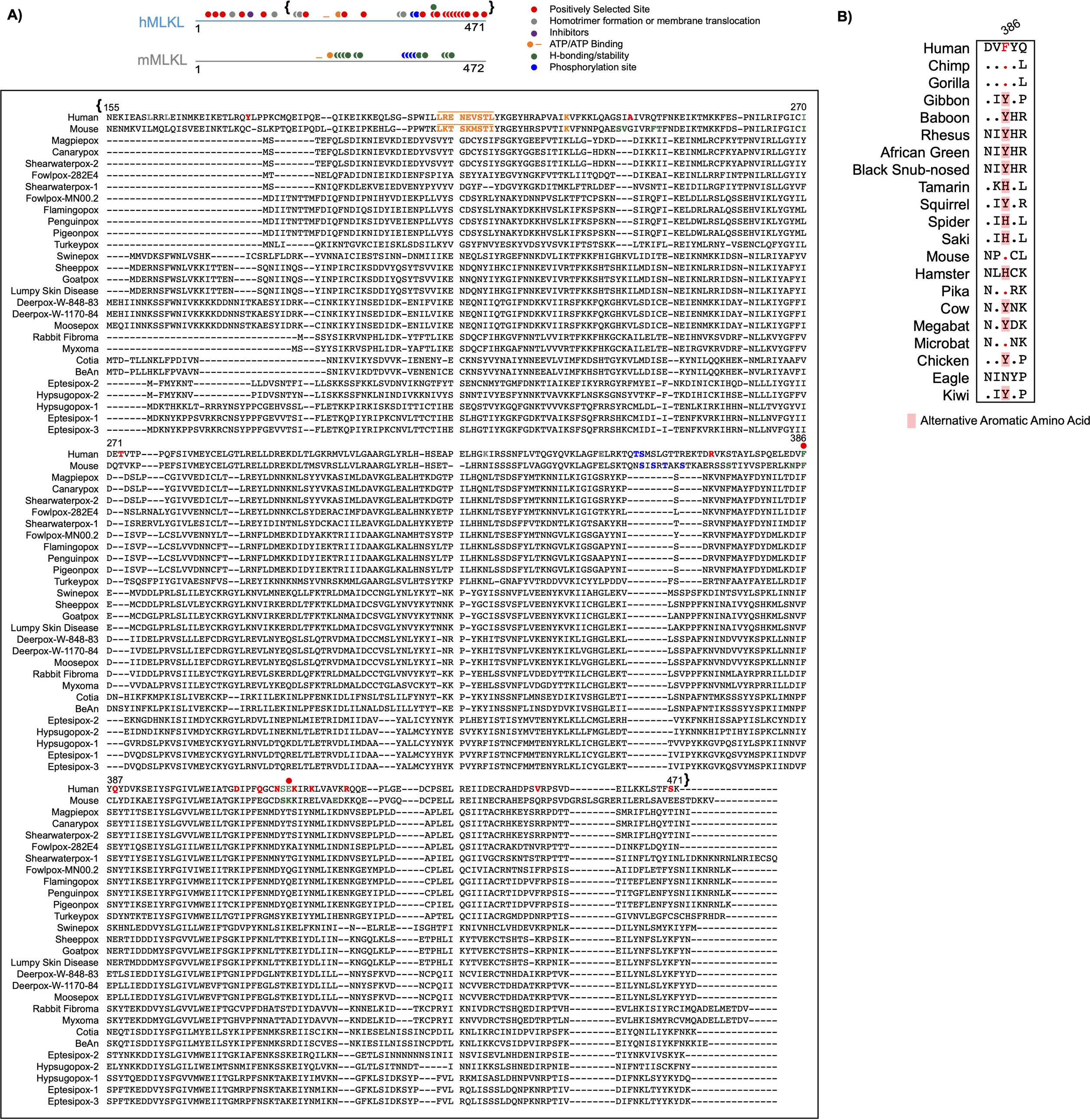
Protein sequence alignments for human MLKL, mouse MLKL and vMLKLs. **A)** Diagrams of hMLKL (light blue) and mMLKL (silver) with previously established functional residues annotated [19-21, 49-51]. Amino acid alignment of hMLKL, mMLKL, and poxvirus MLKLs was constructed using MAFFT, with functional residues annotated on diagrams also highlighted on this alignment. Brackets correspond to the starting point of vMLKL, which is truncated, relative to hMLKL (alignment beginning at hMLKL N155). **B)** Alignment of human MLKL F386 (corresponding to mouse MLKL F373) with MLKLs from diverse host species. Aromatic amino acid positions (red) are primarily conserved to Aves.

**S3 Fig.**
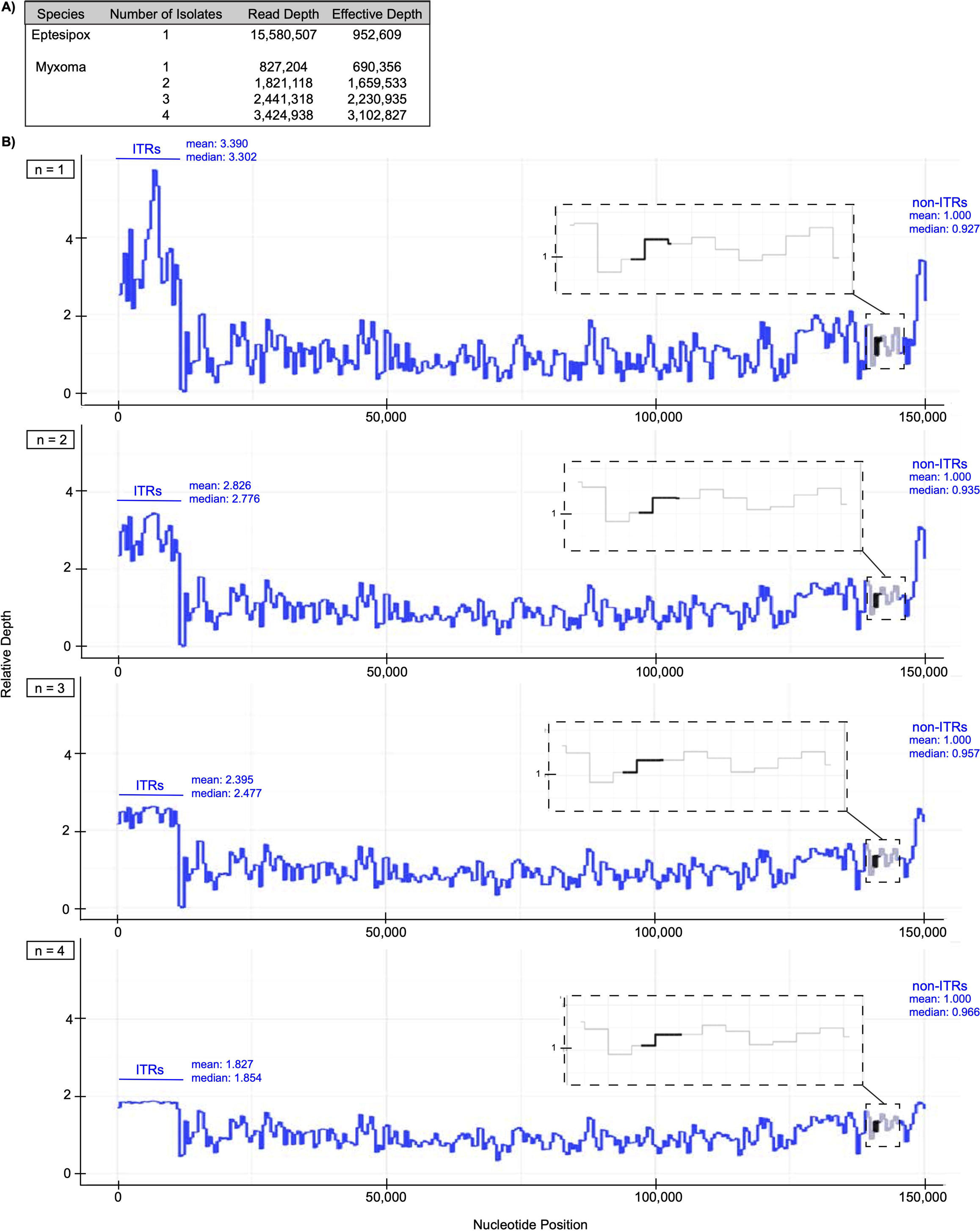
Relative depth analysis of myxoma virus genome sequencing reads indicates a single copy of vMLKL in this species. **A)** Table shows MYXV and EPTV read depth (total number of reads in the BAM file) and effective depth (number of reads in the BAM file that map to the reference genome) following merging of bam files. Analysis performed using data from Kerr *et al.*, and Tu *et al.,* [53, 71]. Myxoma virus analysis was performed as a control given complementary analysis (Fig 4F) and predicted single-copy vMLKL in this and other clade II poxviruses. As the effective depth for myxoma virus sequencing was lower than eptesipoxvirus data, read depth isolates were pooled together to facilitate a more normalized approximation of the read counts per nucleotide in the genome. **B)** Myxoma virus genome relative depth analysis - (reads that align with the ITR and non-ITR regions), where “n =” represents the number of myxoma virus isolates pooled together from Project PRJNA513218 [53]. Inset, relative depth at MYVX vMLKL (black) locus including two flanking upstream and downstream genes (gray). The non-ITR and ITR means and medians are present on each plot.

**S4 Fig.**
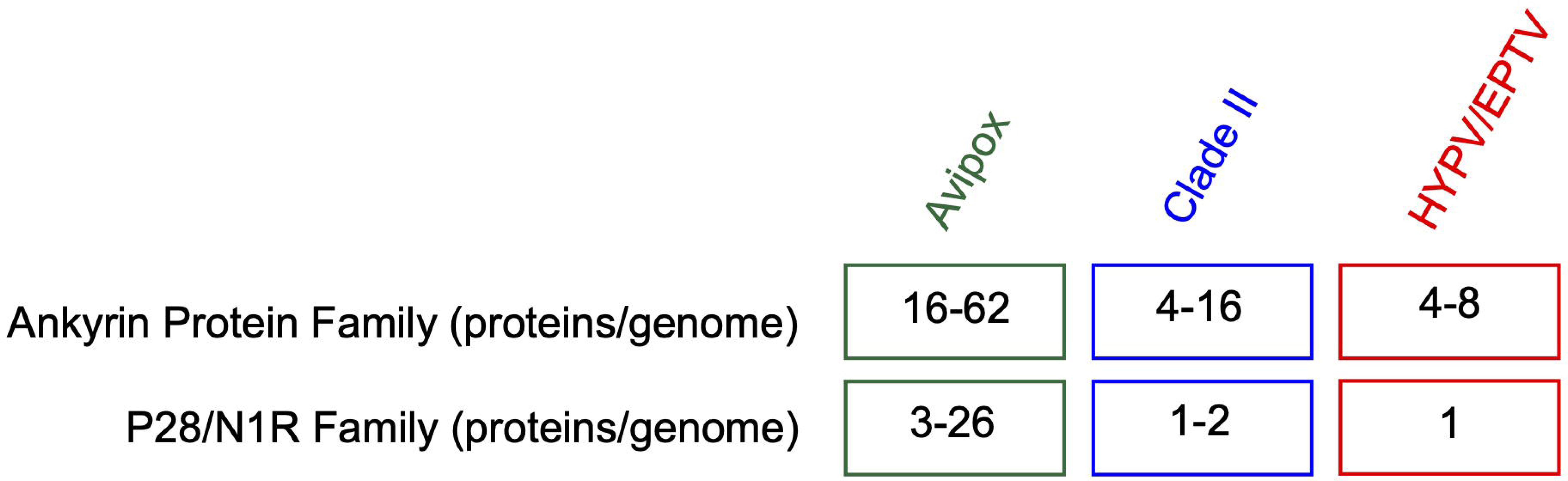
Poxvirus Ankyrin and P28/N1R protein family ranges separated by genus. Summary of ankyrin-repeat containing and P28/N1R proteins per genome analyzed in Fig 4G. Functional protein predictions were not performed for these families.

**S5 Fig.**
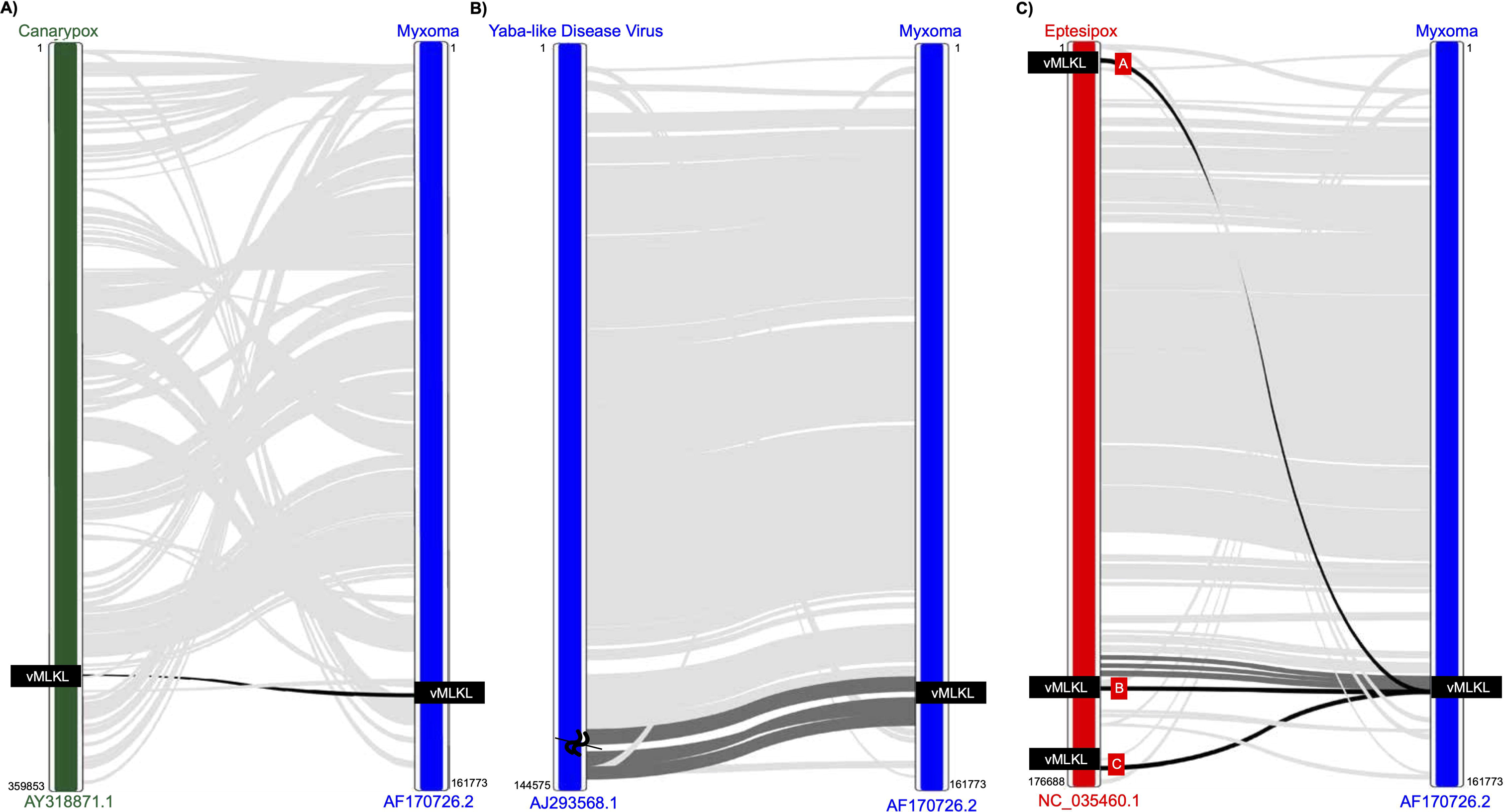
Whole genome synteny analysis illustrates the presence of distinct, multiple copies of vMLKL across poxvirus genomes. The whole genome comparisons are **A)** canarypox virus (*Avipoxvirus* - green) versus myxoma virus (Clade II - blue), **B)** yaba-like disease virus (Clade II - blue) versus myxoma virus and **C)** eptesipoxvirus (Other - red) versus myxoma virus. Sequence accession numbers are centered at the bottom of each genome. Lines that extend between genomes represent homologous regions, that were calculated using CoreGenes 5.0 (https://coregenes.ngrok.io/) [77]. Lines that are light gray represent conserved homologous regions that are not part of the Fig 5 high resolution synteny analysis. Lines in dark gray represent conserved homologous genes that are shown in Fig 5 and the S3 File. vMLKL is represented by a black line. “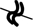” represents a deletion event.

**S1 File. Supporting analysis for Figs 2 and 3 and S1 Fig.**

**S2 File. Supporting analysis for Fig 4 and S4 Fig.**

**S3 File. Supporting analysis for Fig 5.**

**S4 File. Supporting analysis for Figs 6 and 7.**

